# Intricate microbiome differences observed in lactating cows across methane intensity phenotypes

**DOI:** 10.64898/2025.12.11.692445

**Authors:** Adrian Omar Maynez-Perez, Hendra Nur Cahyo, Puchun Niu, Velma T. E. Aho, Phillip B. Pope, Angela Schwarm

## Abstract

Methane emissions from ruminants can be expressed through several metrics as total production, yield, or intensity, each reflecting distinct aspects of energy utilization. Among these, methane intensity defined as grams of methane emitted per kilograms of energy-corrected milk, directly links emissions to productive efficiency; however, the microbial mechanisms underlying variation in this trait remain unclear. Here, we applied genome-resolved metagenomics and metatranscriptomics to characterize rumen microbial identity, functional potential, and transcriptional activity in lactating cows differing in methane intensity while sharing breed and diet. Microbial community composition and diversity were comparable across phenotypes. However, rumen microbial gene expression revealed marked functional divergence. The rumen content of low-methane intensity cows showed enriched transcription of fructan-degrading carbohydrate-active enzymes and butyrate-forming pathways, primarily encoded by *RUG440* (*Atopobiaceae*) and *Sodaliphilus*-affiliated metagenome-assembled genomes. These functions suggest a fructan-butyrate metabolic axis supported by potential cross-feeding between primary degraders and butyrate producers. Conversely, the high-methane intensity rumen exhibited greater transcription of pectin-degrading carbohydrate-active enzymes, mainly carried by *Prevotella* metagenome-assembled genomes, suggesting methyl-ester hydrolysis and methanol release. Despite higher methanogenesis transcript levels in high-methane intensity cows, total methane production did not differ between groups. Together, these findings reveal two contrasting functional configurations of the rumen microbiome in Norwegian Red dairy cattle: a fructan-butyrate-oriented metabolism in low-methane intensity cows and a pectin-methanol-oriented metabolism in high-methane intensity counterparts. This study provides genome-resolved, multi-omic evidence that microbial carbohydrate specialization and fermentation routing contribute to methane intensity phenotypes in dairy cows, offering mechanistic insights for improving ruminant climate efficiency.

## Introduction

The rumen hosts a complex anaerobic microbiome that transforms fibrous plant biomass into metabolites sustaining the ruminant host [1]. This community conformed by bacteria, protozoa, archaea and fungi, acts synergistically to deconstruct plant polysaccharides into intermediate sugars, which are subsequently fermented to short-chain fatty acids (SCFAs). These processes fuel the host and allow conversion of structural carbohydrates into digestible energy [2]. Fermentation inherently produces hydrogen (H_2_), which is mainly consumed by methanogenic archaea to form methane (CH_4_), competing with alternative hydrogenotrophs for this electron source [3, 4]. Thus, the rumen ecosystem regulates both carbon flow from substrates to SCFAs and electron flow through H_2_ sinks [5, 6].

CH_4_ produced in the rumen represents an energy loss of ∼6% of gross energy intake for the animal and accounts for ∼30% of global anthropogenic CH_4_ emissions from livestock [7, 8]. Several metrics are used to evaluate CH_4_ efficiency in dairy cattle, including daily CH_4_ output and CH_4_ yield (g CH_4_ kg⁻¹ dry-matter intake). However, these indicators are not directly linked to production. In contrast, CH_4_ intensity (g CH_4_ kg⁻¹ energy-corrected milk, ECM) is strongly associated with milk output [9], making it a meaningful measure of both feed and production efficiency. Variation in CH_4_ intensity under similar conditions reflects differences in animal energy partitioning, integrating host physiology and microbial energy conversion efficiency.

Rumen energy conversion begins with microbial digestion of plant carbohydrates via carbohydrate-active enzymes (CAZymes), which define microbial niches by targeting distinct substrates [10]. The structure of these niches influences subsequent fermentation routes and redox balance, shaping SCFA profiles and methanogenic fluxes [11, 12]. SCFAs supply ∼70% of the cow’s daily energy, and their composition affects host performance and health. Butyrate fuels the rumen epithelium, whereas acetate and propionate supply carbon for lipogenesis and gluconeogenesis, respectively [13]. During carbohydrate fermentation, SCFA formation also governs hydrogen flow. Acetate and butyrate production release four and two molecules of H_2_, respectively, whereas propionate formation consumes two H_2_ molecules [14]. Therefore, differences in substrate degradation and downstream metabolic routing may underlie CH_4_ intensity phenotypes, influencing both milk production and CH_4_ release.

Advances in next-generation sequencing have expanded understanding of rumen microbiota–CH_4_ relationships. However, studies focused on CH_4_ intensity have largely emphasized taxonomic differences through 16S rRNA gene sequencing [15], while long-read metagenomics research has mainly targeted CH_4_ production or yield [16, 17]. Genome-resolved multi-omics, like integrating metagenomics with metatranscriptomics, enables linking microbial identity, genetic potential, and *in situ* activity, providing a more comprehensive view of microbial metabolism [18]. Although multi-omics has been applied to animals differing in CH_4_ intensity [19], these studies typically compared distant breeds to capture large-effect differences. To our knowledge, a genome-resolved multi-omic analysis of CH_4_ intensity phenotypes in genetically similar animals, allowing the resolution of subtle yet functionally meaningful distinctions, has not yet been reported.

Here, we applied genome-resolved metagenomics and metatranscriptomics to link rumen microbial identity, functional potential, and gene expression in cows differing in CH_4_ intensity while sharing breed and diet. We specifically assessed whether transcriptional activity of CAZymes and central fermentation pathways diverged between phenotypes. Our results reveal distinct functional configurations between low- and high- CH_4_ intensity cows, characterized by contrasting carbohydrate-degradation preferences and downstream metabolic routing.

## Material and Methods

See Supplementary Materials and Methods for further details.

### Experimental design and CH_₄_ intensity classification

The animal trial was conducted at the Norwegian University of Life Sciences (Ås, Norway) under approval from the Norwegian Food Safety Authority (Mattilsynet; FOTS ID 29827). Total of 16 lactating Norwegian Red cows assigned into low- (LMI) and high-CH₄ intensity (HMI) groups (n = 8 per group) calculated from daily CH_4_ production and energy-corrected milk (ECM) output (CH_4_/ECM) were enrolled in a 17-day continuous experiment consisting of an 11-day adaptation and 6-day sampling period. All cows were fed grass silage and concentrate (83:17 ratio on DM basis during the experiment) with similar nutrient composition (Table S1) and maintained under uniform management conditions. Cows averaged 251 ± 30 days in milk, 638 ± 86 kg body weight, and 18 ± 4 kg milk yield at the start of the experiment.

### Feed intake, milk yield, and gas emission measurements

Individual dry matter intake (DMI) was calculated daily as feed offered minus refusals. Milk yield was recorded twice daily, and ECM was estimated according to standard energy-correction equations. CH_4_ and carbon dioxide (CO_2_) emissions were measured using a GreenFeed system (C-Lock Inc., USA) over six consecutive days, with measurements staggered 5 to 6-hour intervals (22 times per cow) spanning nearly the full 24-hour cycle. Each cow was sampled for ∼5 min per session while receiving small concentrate baits. Background air was recorded between measurements, and gas concentrations were automatically corrected by the GreenFeed system. Calibration procedures and recovery tests are detailed in supplementary methods.

### Sample collection and processing

Rumen fluid was collected from all cows on day 17 using an esophageal tubing system (Selekt™, Nimrod Veterinary Products Ltd., UK). To minimize saliva contamination, the initial ∼200 mL of aspirated content was discarded. Subsequently, approximately 200 mL of rumen fluid was collected from each cow, from which 2 mL were subsampled into sterile cryovials, snap-frozen in liquid nitrogen, and stored at −80 °C until microbial analyses.

### Metagenomic analysis

DNA extraction and sequencing as well as initial metagenomic sequence data analysis for rumen digesta samples was performed at DNASense ApS (Aalborg, Denmark).

#### DNA extraction and sequencing

Rumen fluid samples were processed for microbial DNA extraction using the DNeasy PowerSoil Pro Kit (Qiagen, Germany) following the manufacturer’s protocol, with a mild horizontal bead-beating step of 10 min at maximum speed. DNA extracts were further purified using the DNeasy PowerClean Pro Cleanup Kit (Qiagen). DNA concentration and purity were assessed with a Qubit dsDNA HS Assay Kit and a NanoDrop One spectrophotometer (Thermo Fisher Scientific, USA), while fragment size distribution was evaluated using Genomic DNA ScreenTapes on an Agilent TapeStation 4200 (Agilent Technologies, USA).

Barcoded DNA libraries were prepared using the SQK-LSK114.24 ligation sequencing kit (Oxford Nanopore Technologies, UK) with minor protocol modifications. Approximately 10–20 fmol of pooled, barcoded DNA were loaded onto a primed FLO-PRO114M (R10.4.1) flow cell and sequenced on a PromethION P2 Solo device running MinKNOW v22.12.5. Basecalling and demultiplexing of FAST5 signal data were performed using Guppy v6.4.6 (Oxford Nanopore Technologies) with the super-accurate model (dna_r10.4.1_e8.2_400bps_sup.cfg) and midstrand detection enabled.

#### Quality control and assembly

Raw reads were quality filtered, and sequencing statistics were summarized using Nanoq v0.10.0 [20]. A draft de novo co-assembly was generated using Flye v2.9.1b1780 [21] with approximately 7.5 Gbp of data per sample. The assembly was polished once using Medaka v1.7.2 (Oxford Nanopore Technologies). Assembly graphs were visualized with Bandage v0.8.1 [22], and contigs shorter than 1,000 bp were removed using SeqKit v2.2.0 [23].

#### Metagenome binning and refinement

Automated binning was performed using MetaBAT2 v2.15 [24] and Vamb v4.1.3 [25]. The completeness and contamination of metagenome-assembled genomes (MAGs) were evaluated with CheckM2 v1.0.1 [26, 27]. MAG dereplication and quality filtering were conducted using dRep v3.4.3 [28] with the following thresholds: minimum MAG length 0.5 Mbp, completeness ≥1%, and contamination ≤10%. A total of 1,157 MAGs were recovered, of which 424 passed MIMAG quality criteria [29]: high-quality (HQ, ≥90% completeness and ≤5% contamination) and medium-quality (MQ, ≥50% completeness and ≤10% contamination). For comparative analyses between CH_4_ intensity phenotypes, MAGs present in fewer than one-third of the samples were removed (n=6). Abundance data were subsequently normalized to relative abundance per sample for downstream statistical analyses.

#### Taxonomic and functional annotation

MAG taxonomy was defined with GTDB release r214 using GTDB-Tk v2.3.0 [30]. Functional annotation of translated amino acid sequences was performed with DRAM v1.4 [31] using UniRef as reference (DRAM.py annotate_genes --use_uniref --threads 64). Annotated features from DRAM were used to interpret metabolic potential and to integrate with metatranscriptomic results. The presence of carbohydrate-active enzymes (CAZymes) was determined with dbCAN v4.0.0 [32], combining HMMER, dbCAN-sub, and DIAMOND searches against the CAZy database. Only CAZymes supported by all three methods were retained for downstream analyses.

### Metatranscriptomic analysis

RNA extraction, sequencing and mapping of the sequences against reference databases were performed at DNASense ApS (Aalborg, Denmark).

#### RNA extraction and sequencing

Total RNA was extracted from rumen content using the RNeasy PowerMicrobiome Kit (Qiagen, Germany) following the manufacturer’s protocol with minor modifications. Specifically, custom reagent volumes were applied, and bead beating was performed at 6 m/s for 4 × 40 s. RNA integrity and purity were assessed by gel electrophoresis on a TapeStation 2200 using RNA ScreenTape (Agilent Technologies, USA). RNA concentration was quantified with the Qubit RNA HS/BR Assay Kit (Thermo Fisher Scientific, USA). Residual DNA was removed using the TURBO DNA-free Kit (Thermo Fisher Scientific, USA), followed by an additional quality control step using TapeStation and Qubit assays. Ribosomal RNA was depleted with the Ribo-Zero Plus rRNA Depletion Kit (Illumina, USA), and residual DNA was further eliminated using the DNase MAX Kit (MoBio Laboratories, USA). The resulting RNA was purified with CleanPCR SPRI beads (CleanNA, Netherlands) and used for library construction with the NEBNext Ultra II Directional RNA Library Preparation Kit (New England Biolabs, USA). Library concentration was determined using the Qubit HS DNA Assay, and fragment size was verified on a TapeStation with D1000 ScreenTape. Libraries were pooled in equimolar concentrations and sequenced on an Illumina NovaSeq platform (2 × 150 bp paired-end reads). All protocols were followed according to manufacturer recommendations with minor modifications. Metatranscriptomic sequencing was successful for 15 of the 16 rumen samples (HMI n = 8; LMI n = 7); one library failed quality control and was excluded from downstream analyses.

#### Read processing and mapping to MAGs

Raw metatranscriptomic reads were mapped against the *Bos taurus* reference genome (ARS-UCD1.3) using minimap2 v2.2 to remove host-derived sequences. Non-paired mapped reads were extracted with samtools v1.17 using parameters samtools fastq -f12 -F256 -c7 −1 read1.fq.gz −2 read2.fq.gz. Remaining rRNA sequences were removed in silico with SortMeRNA v4.3.6 [33] using SILVA reference databases (silva-bac-16s-id90, silva-arc-16s-id95, silva-bac-23s-id98, silva-arc-23s-id98, silva-euk-18s-id95, silva-euk-28s-id98) and parameters --out2 --paired_out --fastx --threads 64. Cleaned reads were pseudo-aligned and quantified against the rumen microbial genome catalog (MAGs) using Kallisto v0.50.0 [34]. The resulting transcript-level abundance tables were imported and summarized to the gene level in R (v4.2.2) using the tximport v1.26.1 package [35].

#### Transcript quantification and normalization

Genes with low expression were filtered out to reduce noise, retaining only those with at least one count per million (CPM) in 20% or more of the samples. To correct for differences in sequencing depth across libraries, normalization factors were calculated using the Trimmed Mean of M-values (TMM) method implemented in edgeR [36]. Normalized data were subsequently transformed to log₂ counts per million (log-CPM) for differential expression and gene set analyses, and to log₁₀ transcripts per million (log₁₀ TPM) for visualization of functional profiles at the MAG or gene level. For exploratory analyses and quality control, variance-stabilizing transformation (VST) values were also generated using DESeq2 [37], ensuring consistent variance across the expression range.

### Statistical analysis

Statistical analyses were performed in R (v4.3.2). For animal performance data, differences between low- and high– CH_4_ intensity (LMI and HMI) cows were evaluated using linear mixed-effects models implemented in the nlme package [38]. The model included CH_4_ intensity group, parity, and their interaction as fixed effects, and individual cow as a random effect. Least-square means were estimated and P-values compared using Benjamini–Hochberg.

For metagenomic data, α-diversity indices (richness, Shannon, Simpson) were computed using the vegan package [39] and compared between groups using linear models with CH_4_ intensity group and parity as fixed effects. β-diversity was assessed using Bray–Curtis dissimilarity based on relative MAG abundance profiles, visualized by principal coordinates analysis (PCoA). Group differences were tested with PERMANOVA (adonis2, 999 permutations), and homogeneity of multivariate dispersion was verified with betadisper. Relative abundances of MAGs associated with differential carbohydrate-degrading and/or metabolic pathways (described below) were compared between phenotypes using Wilcoxon rank-sum tests. For co-abundance relationships among these MAGs Pairwise Spearman correlations (ρ) were computed across samples using the Hmisc package [40].

For metatranscriptomic data, sample-wise variance stabilization was performed using DESeq2. Ordination of global expression profiles was evaluated by principal component analysis (PCA) on variance-stabilized counts, and statistical separation between groups was tested with PERMANOVA using Euclidean distances. At the functional level, transcripts were categorized into metabolic pathways and carbohydrate-degrading enzymes based on their KO and CAZyme family respectively (Supplementary File S1). Gene sets were assessed using the limma–voom framework applied to log₂ counts-per-million (CPM). Gene set enrichment was tested with the CAMERA method [41] using the limma package [42]. All statistical analyses were corrected using the Benjamini–Hochberg false discovery rate (FDR). Significance thresholds were set at FDR□<□0.05 and tendency at 0.05 ≤ FDR□<□0.10.

## Results

### Differential energy partitioning to milk across divergent CH_₄_ intensity phenotypes

We classified 16 cows with comparable body weight, days in milk, and diet (Fig. 1B) into low- and high- CH_4_ intensity groups (LMI and HMI; n=8 each; Fig. 1A). LMI cows produced more energy-corrected milk (ECM; FDR < 0.01), showed higher ECM/DMI (FDR < 0.01), and had lower CH_4_ intensity (CH_4_/ECM; FDR < 0.01) than HMI cows, whereas DMI, dry-matter digestibility (DMD), and CH_4_ production did not differ between groups (Fig. 1B). Milk composition (fat, protein and lactose) was similar across phenotypes (Table S2). Total energy intake (gross, digestible and metabolizable) was also comparable, however LMI cows deposited more energy in milk (MJ d⁻¹; FDR < 0.01), resulting in a higher conversion of gross energy to milk energy (17.8% in LMI vs 14.0% in HMI; FDR < 0.01; Table S2). CH_4_ and CO_2_ yields normalized by body weight or DMI were not different, whereas their intensities per ECM were lower in LMI (CH_4_: 20.7 vs 27.6 g kg⁻¹ ECM; FDR < 0.01; CO_2_: 587 vs 794 g kg⁻¹ ECM; FDR < 0.01; Table S2). The *in vivo* experiment demonstrated that, despite similar feed intake, digestibility, and emissions across phenotypes, LMI cows partitioned a greater fraction of available energy into milk.

**Figure 1.**
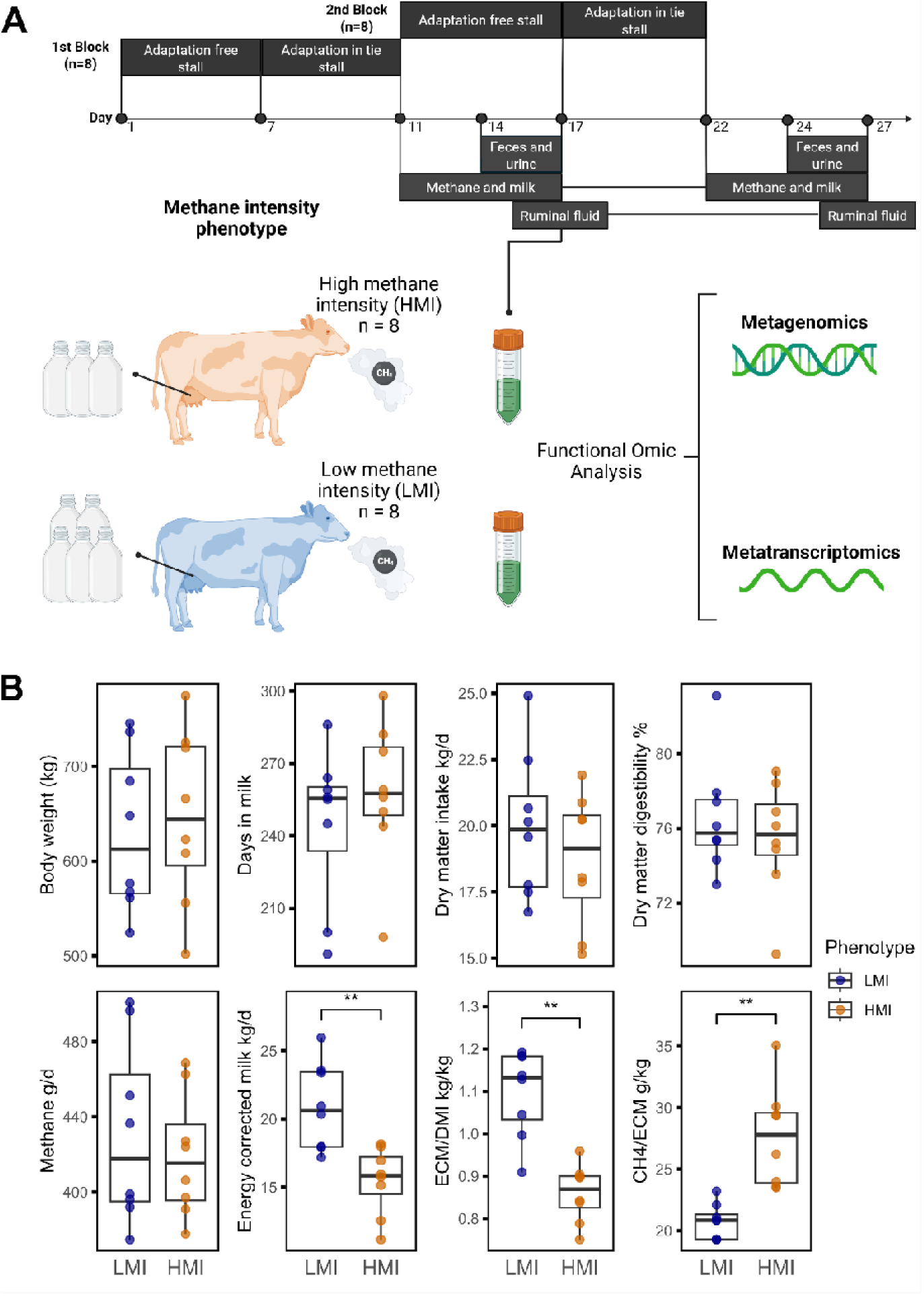
Experimental design and host characteristics. **A.** Experimental design. A total of 16 Norwegian Red dairy cows, selected based on CH_4_ intensity (LMI: Low CH_4_ intensity; HMI: High CH_4_ intensity), were divided into two blocks (n=8 per block). The experiment consisted of an adaptation period (7-day in a free stall, followed by 4 days in a tie stall), and a 5-day sampling period, during which faeces, urine, CH_4_, milk, and ruminal fluid were collected. A functional multi-omic analysis, including metagenomic and metatranscriptomic approaches, was performed on the ruminal fluid samples. **B.** Host characteristics. Boxplot graphs show the results of both phenotypes for the following parameters: body weight (kg), days in milk (d), dry matter intake (kg/d), dry matter digestibility (%), CH_4_ production (g/d), energy-corrected milk (kg/d), ECM/DMI (kg/kg), and CH_4_/ECM (g/kg). Statistical significance was determined using a linear mixed-effects model (adjusted by the Benjamini–Hochberg method), with significant P-values indicated as follows: *FDR < 0.05, **FDR < 0.01, ***FDR < 0.001. Created in BioRender. Maynez Perez, A. (https://BioRender.com/mgynu71).

### Rumen microbiome composition is similar across CH_₄_ intensity phenotypes

We recovered 424 genome-resolved populations (MAGs) meeting MIMAG criteria (73 high-quality; 351 medium-quality), with a mean completeness of 75.5 ± 14.5% and contamination of 2.17 ± 2.14% (mean genome size 2.26 Mb; Fig. 2A). Sixty-one percent were classified to species level. On average, 16.83% of reads mapped to MAG contigs (Fig. 2A). Using the Genome Taxonomy Database (GTDB), the catalogue comprised 24 archaeal and 400 bacterial MAGs. Dominant phyla across both phenotypes included *Bacteroidota*, *Bacillota*, *Methanobacteriota*, *Fibrobacterota*, *Actinomycetota*, and *Pseudomonadota*; prevalent families included *Bacteroidaceae*, *Lachnospiraceae*, *Methanobacteriaceae*, *Fibrobacteraceae*, CAG-74, *Atopobiaceae*, UBA932, *Oscillospiraceae*, *Paludibacteraceae*, and *Ruminococcaceae*. Common genera were *Prevotella*, *Fibrobacter*, *Methanobrevibacter_A*, CAG-791, RUG740, RUG440, *Cryptobacteroides*, RF16, *Sodaliphilus*, and *Limivicinus* (Fig. S1).

**Figure 2.**
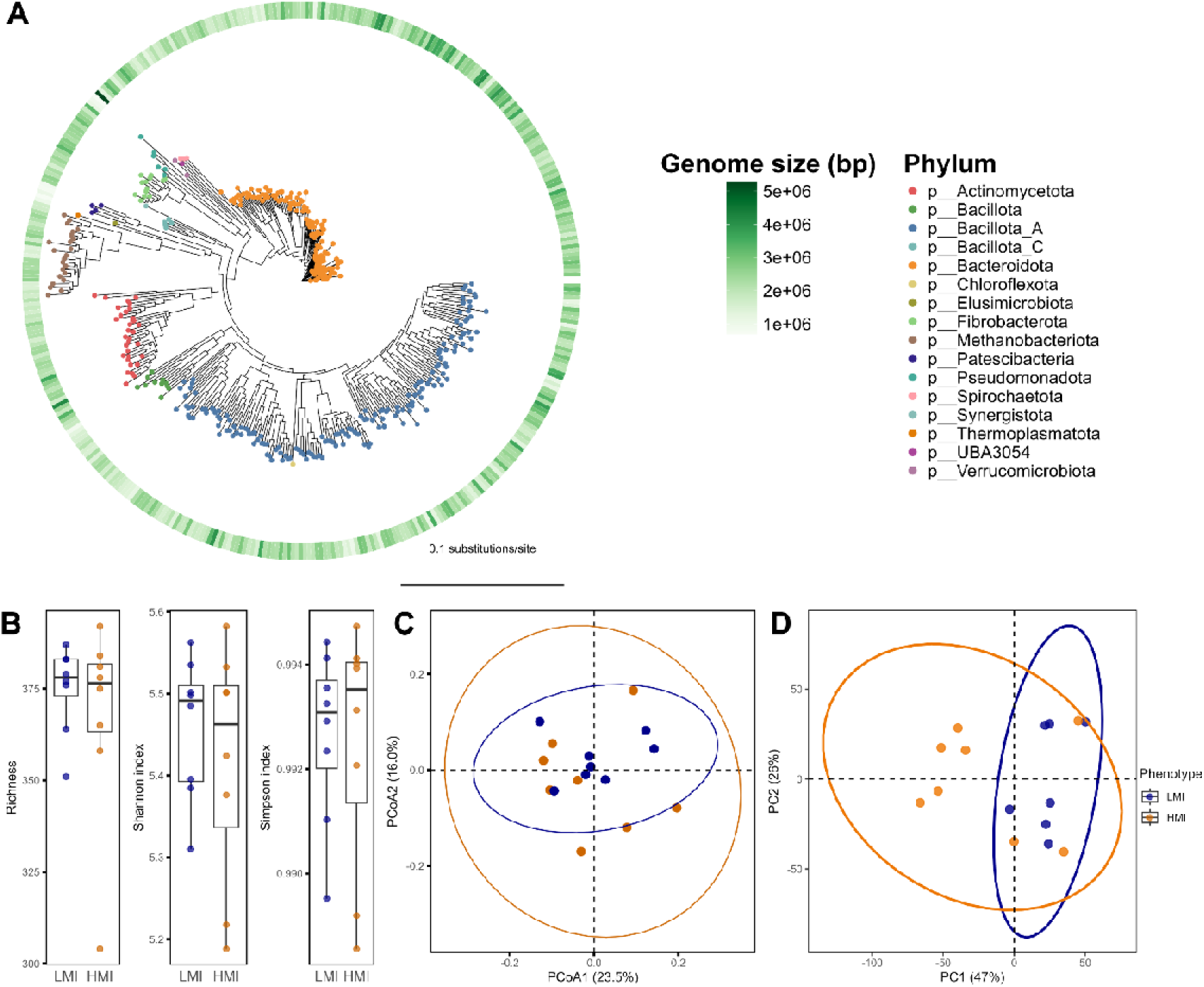
Different CH_4_ intensity phenotypes show similar microbial structure. **A.** The maximum-likelihood tree with 424 high and medium quality MAGs; clades are colored according to the taxonomic classification of genomes at phylum level; the heatmap in the outer layer shows genome size corresponding to each genome. **B.** Boxplots displaying alpha diversity including richness, Shannon and Simpson index. No difference was found in any of the metrics (P > 0.05). **C.** Principal coordinates analysis (PCoA) plot illustrating beta diversity based on Bray-Curtis method. No difference was found (P > 0.05). **D.** Principal component analysis (PCA) of gene expression profiles in ruminal content of low (LMI) and high (HMI) CH_4_ intensity cows. PCA was conducted using variance-stabilizing transformation (VST) normalized counts from DESeq2. No difference was found (P > 0.05).

In analyses of microbial diversity based on metagenome abundances, alpha-diversity (richness, Shannon, Simpson) did not differ between LMI and HMI (LMM; FDR = 0.86, 0.86, 0.97; Fig. 2C). Beta-diversity based on Bray–Curtis dissimilarity also showed no separation by phenotype (PERMANOVA; 1000 permutations: FDR = 0.22; Fig. 2D), with no evidence of heterogeneous dispersion (betadisper: FDR = 0.26). Analysis of the unbinned metagenomic fraction at species level likewise showed no differences in α- or β-diversity between groups (Fig. S2). Considering the metatranscriptomic data, 1,749,857 transcripts were successfully mapped to MAGs, with a mean of 9,341,671 ± 1,408,685 transcripts per sample; library sizes after DESeq2 variance-stabilizing transformation (VST) were comparable between phenotypes (FDR = 0.15; Fig. 2D), with no evidence of heterogeneous dispersion (betadisper: FDR = 0.23). These data indicate community composition is broadly similar across divergent CH_4_ intensity phenotypes.

### Functional activity differentiates CH_4_ intensity phenotypes

We observed consistent differences in carbohydrate utilization and electron flow between the two contrasting CH_4_ intensity phenotypes. To compare gene-level activity of carbohydrate degradation and fermentation pathways, we analysed transcriptional differences between the microbiomes of the two phenotypes and tested predefined gene sets with CAMERA, which accounts for inter-gene correlation. Each substrate or pathway was then summarized by the distribution of moderated t-statistics across its member genes (Fig. 3). Among the total number of CAZymes detected using metatranscriptomics across our data (Fig. 3A), those putatively active on fructan (183 genes) were enriched in LMI (FDR = 0.017), whereas pectin-degrading CAZymes (400 genes) showed higher expression in HMI (FDR = 0.03). CAZymes targeting cellulose, hemicellulose, xylan, starch, arabinan, and chitin were not significant after FDR correction (all FDR ≥ 0.52). For central metabolic pathways (Fig. 3B), transcripts mapping to the butyrate pathway (189 genes) were enriched in LMI (FDR = 0.002), while acetate, propionate (succinate or acrylate routes), lactate, ethanol, and Wood–Ljungdahl modules did not differ between phenotypes (all FDR ≥ 0.18). Collectively, these results indicate a functional axis in LMI cows characterized by greater transcription of fructan-degrading CAZymes and butyrate-forming enzymes, contrasted with a pectin-degrading signature in HMI cows.

**Figure 3.**
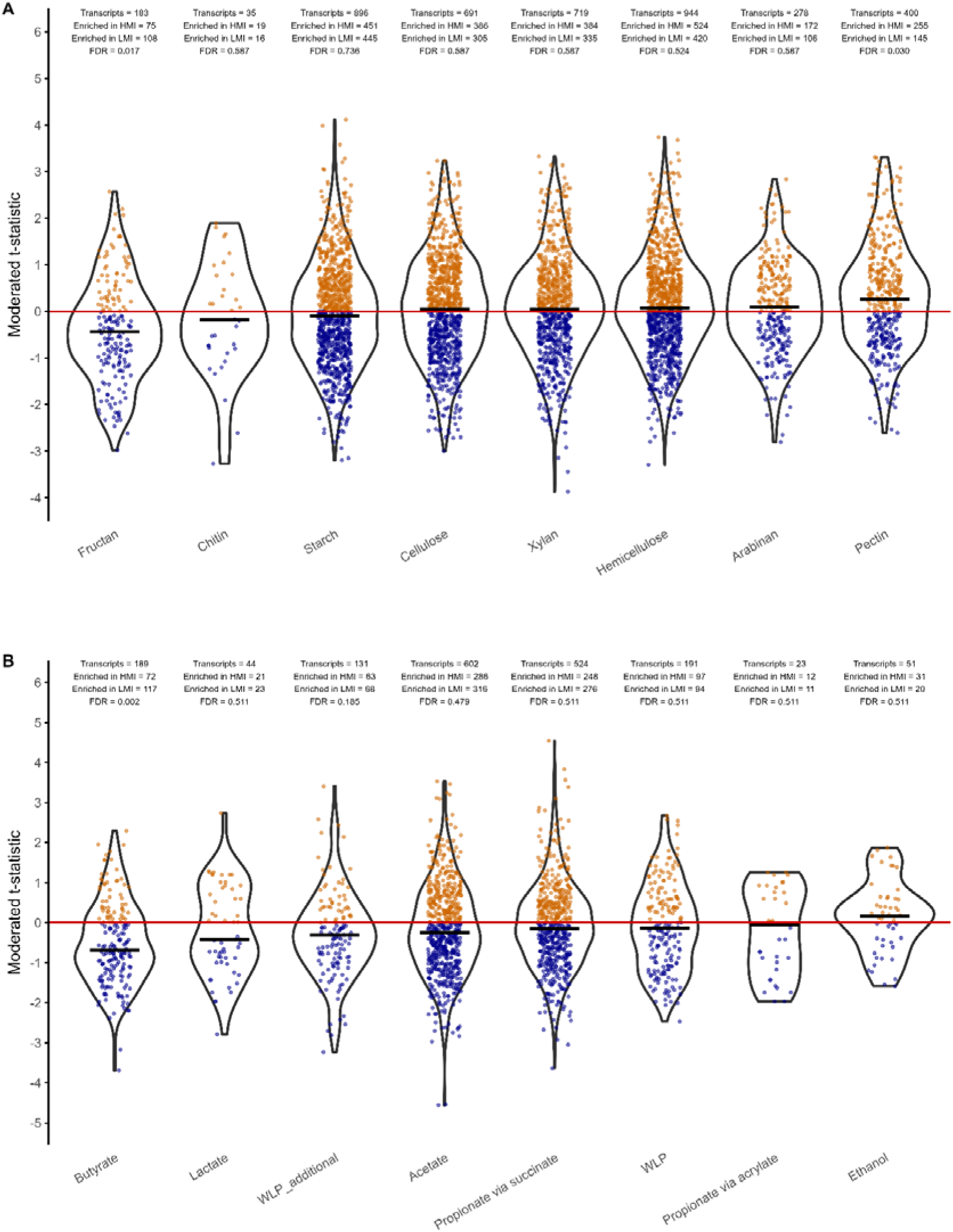
Differences in ruminal metabolism and carbohydrate degradation between CH4 intensity groups. **A.** Comparative metatranscriptomic analysis of rumen carbohydrate-active enzymes (CAZymes) grouped by substrate. **B.** Comparative metatranscriptomic analysis of rumen metabolic pathways (KEGG). Each dot (gene) represents the moderated t-statistic from a limma–voom model contrasting phenotypes (HMI = high CH4 intensity vs LMI = low CH4 intensity). Outlines depict the kernel density of gene-level t-statistics per set, whereas the black crossbar marks the median. The colour of each dot encodes direction where positive values (orange) indicate higher expression in the HMI group, whereas negative values (blue) indicate higher expression in the LMI group. Statistical significance was determined by CAMERA gene-set testing applied to the voom (log-CPM) matrix, adjusted by FDR (Benjamini–Hochberg).

We further examined the methanogenesis modules to assess whether these metabolic shifts affected the terminal steps of CH_4_ formation. All four gene sets including CO_2_ to CH_4_, methanol to CH_4_, shared methanogenesis, and coenzyme M formation showed positive moderated t-statistics, indicating higher transcript abundance in HMI cows (Fig. S3). However, none reached statistical significance after FDR correction (all FDR ≥ 0.60). This suggests that while HMI rumen communities transcribe methanogenesis genes more intensely overall, the magnitude of difference is small compared to the pronounced transcriptional divergence in carbohydrate degradation and fermentation pathways. These observations were within expectations given we observed no difference in absolute CH_4_ measurements across the CH_4_ intensity phenotypes.

### Fructan-targeting CAZymes co-occur with enrichment of butyrate pathway in LMI rumen

To identify which specific microbial populations encoded the fructan-degrading CAZymes enriched in LMI cows (FDR = 0.017; Fig. 4A), we compared metagenomic abundances of MAGs containing such enzymes. Wilcoxon tests highlighted six MAGs (MAG_68, MAG_44, MAG_98, MAG_88, MAG_353, MAG_362) which tended to be more abundant in LMI (Fig. 4B; Wilcoxon test, unadjusted P < 0.05). A MAG-by-KO heatmap of GH32 family members (fructan-active CAZymes) confirmed higher transcript levels (log10 TPM) in LMI compared to HMI across these MAGs (Fig. 4C).

**Figure 4.**
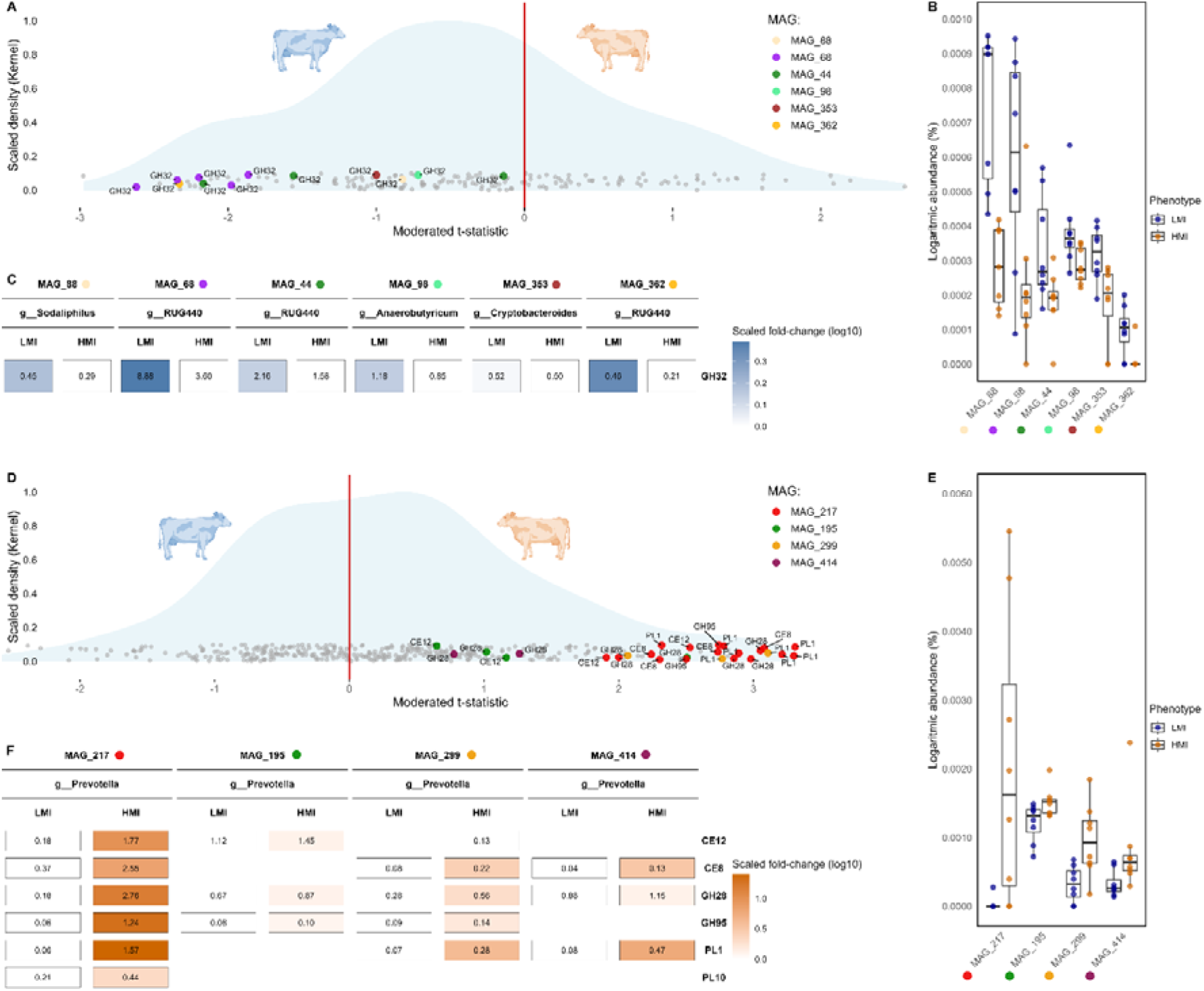
Differential carbohydrate degradation across CH_4_ intensity phenotypes. **A.** MAGs involved in fructan degradation. The blue area represents the kernel density of gene-level t-statistics per set related to fructan degradation. Genes with values above zero indicate higher expression in the HMI group, whereas genes below zero have higher expression in the LMI group. The colour of the dots represents the MAG from which the gene originates, with only genes below zero being coloured. **B.** The box plot displays the metagenomic logarithmic abundance of the MAGs represented in the distribution plot. All the coloured MAGs showed a tendency (non-adjusted P < 0.05 from a two-sided Wilcoxon rank-sum test). **C.** The heatmap illustrates the TPM of fructan degradation CAZymes on trend MAGs in the LMI and HMI phenotypes, with the colour intensity based on the scaled fold-change (log10) against is counterpart on the same MAG, the value inside represents the TPM. **D.** MAGs involved in pectin degradation. The blue area represents the kernel density of gene-level t-statistics per set related to pectin degradation. Genes with values above zero indicate higher expression in the HMI group, whereas genes below zero have higher expression in the LMI group. The colour of the dots represents the MAG from which the gene originates, with only genes below zero being coloured. **E.** The box plot displays the metagenomic logarithmic abundance of the MAGs represented in the distribution plot. All the coloured MAGs showed a tendency (non-adjusted P < 0.05 from a two-sided Wilcoxon rank-sum test). **F.** The heatmap illustrates the TPM of pectin degradation CAZymes on trend MAGs in the LMI and HMI phenotypes, with the colour intensity based on the scaled fold-change (log10) against is counterpart on the same MAG, the value inside represents the TPM.

Moreover, for the butyrate pathway gene set which was also enriched in LMI cows (FDR = 0.002; Fig. 5A), the genes driving this route from acetyl-CoA to butyrate (hbd [K00074], bcd [K00248], atoB [K00626], buk [K00929], atoD [K01034], atoA [K01035], crt [K01715]) were mainly encoded by fructan-degrading populations MAG_98 and MAG_88 as well as MAG_233 and MAG_379, all of which also tended to be more abundant in LMI (Fig. 5B; Wilcoxon test, unadjusted P < 0.05). Similarly, a KO-level heatmap showed numerically greater transcript levels for multiple butyrogenic steps in LMI, particularly for MAG_88, MAG_98 and MAG_233 (Fig. 5C).

**Figure 5.**
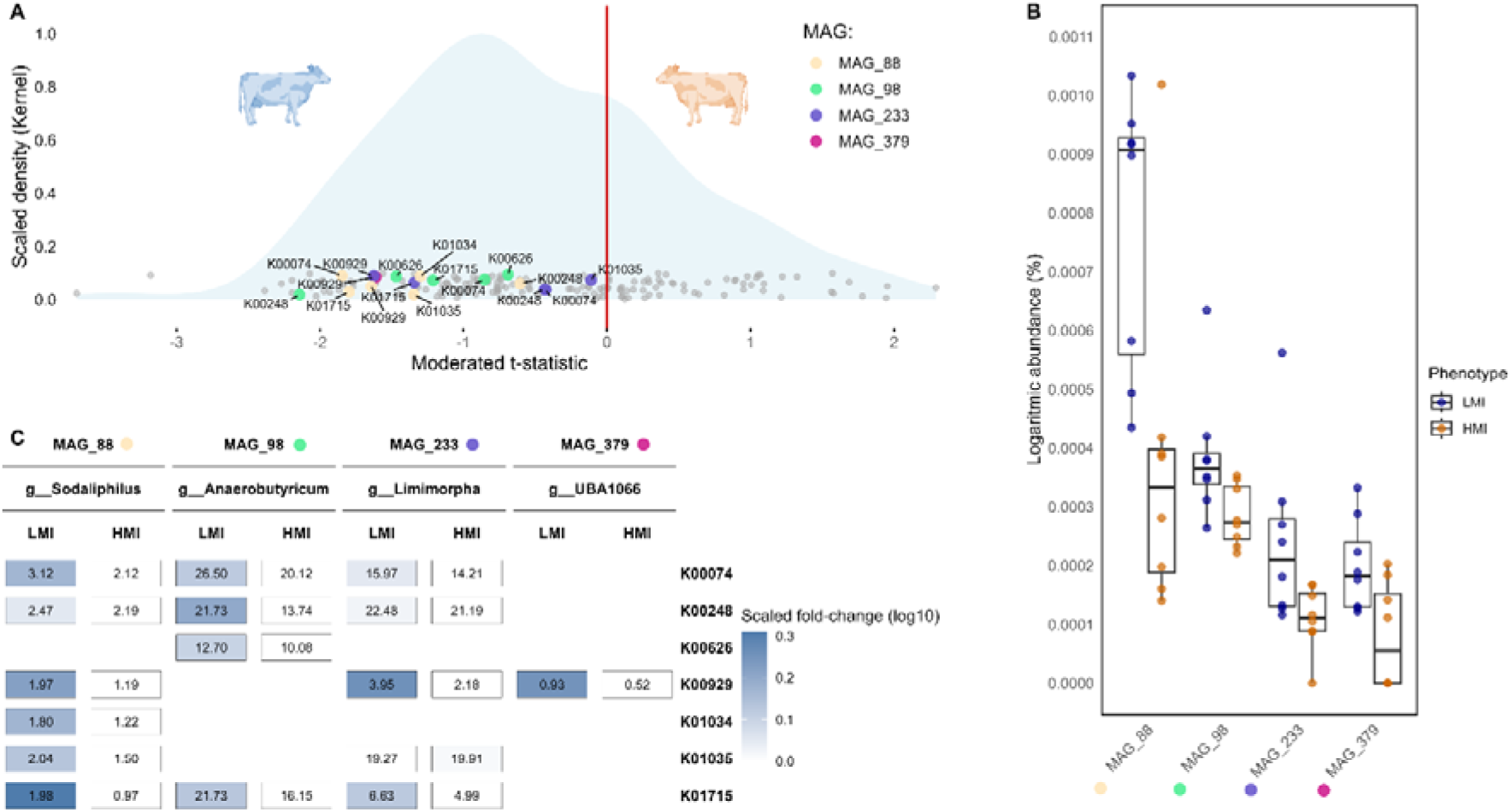
Differential butyrate metabolism across CH_4_ intensity phenotypes. **A.** MAGs involved in butyrate metabolism. The blue area represents the kernel density of gene-level t-statistics per set related to butyrate metabolism. Genes with values above zero indicate higher expression in the HMI group, whereas genes below zero have higher expression in the LMI group. The colour of the dots represents the MAG from which the gene originates, with only genes over zero being coloured. **B.** The box plot displays the metagenomic logarithmic abundance of the MAGs represented in the distribution plot. All the coloured MAGs showed a tendency (non-adjusted P < 0.05 from a two-sided Wilcoxon rank-sum test). **C.** The heatmap illustrates the TPM of butyrate metabolism KOs on trend MAGs in the LMI and HMI phenotypes, with the colour intensity based on the scaled fold-change (log10) against is counterpart on the same MAG, the value inside represents the TPM.

Consistent with these transcriptional patterns, correlation analysis between MAGs and functional traits revealed a positive association between butyrate and fructan modules, which clustered together with LMI-enriched MAGs (MAG_68, MAG_233, MAG_98) and in opposition to pectin-associated taxa (Fig. S4). Genomic trait profiling further confirmed that these LMI-associated MAGs encoded a high abundance of SCFA biosynthesis and polysaccharide degradation traits, particularly linked to poly- and oligosaccharide utilization, sugar transport, and butyrate formation (Fig. S5). Together, these results indicate a coordinated fructan→butyrate axis in the low-CH_4_ intensity rumen, where specific MAG guilds connect carbohydrate specialization with fermentation product profiles.

### Pectin-targeting CAZymes are enriched in the HMI rumen microbiome

The key populations enriched in the HMI rumen microbiome that encoded pectin-degrading CAZymes (FDR = 0.030; Fig. 4D) were affiliated with four MAGs classified as genus *Prevotella* (MAG_217, MAG_195, MAG_299, MAG_414). Their pectinase profile included CAZymes active on homogalacturonan, rhamnogalacturonan I or rhamnogalacturonan II (e.g. GH28, GH95, PL1, PL10) as well as putative methyl-esterases (e.g. CE8) that remove methyl groups from galacturonic acid residues. These same MAGs tended to be more abundant in HMI cows at the metagenome level (Fig. 4E; Wilcoxon test, unadjusted P < 0.05) and showed higher transcription across multiple pectinolytic families (Fig. 4F). Wider correlation analysis grouped pectin-degrading MAGs into the HMI cluster, separated from the LMI cluster of fructan–butyrate-associated taxa (Fig. S4). Functional genome profiling of HMI *Prevotella* MAGs highlighted methyl-group transfer and methanol-linked traits, compatible with cross-feeding to methylotrophic archaea (Fig. S5). Thus, the rumen microbiome of HMI Norwegian red dairy cattle analysed herein favours a pectin-degrading, methyl-releasing niche in contrast to the putative fructan catabolism and butyrate formation observed in the LMI rumen microbiome.

## Discussion

Although overall microbial community structure was similar between cows differing in CH_4_ intensity, functional activity related to carbohydrate degradation and central metabolic pathways differed for several defined concise niches between phenotypes. The LMI rumen microbiome showed higher expression of CAZymes targeting fructan digestion and of genes involved in butyrate metabolism, carried by MAGs that were positively correlated between them (Fig. 6). In contrast, the rumen microbiome of HMI cattle exhibited greater transcription of pectin-degrading enzymes, mainly encoded by *Prevotella*-affiliated MAGs (Fig. 6). Only numerical differences were observed for methanogenesis-related transcripts, and CH_4_ production did not differ at the animal level. Similar patterns where cows producing comparable amounts of CH_4_ differed in CH_4_-production categories because of variation in feed intake or milk output have been reported previously [43, 44]. Genetic differences between CH_4_ intensity groups may contribute to the partitioning of energy into milk, but could also influence microbial dynamics and metabolism, underscoring a microbial signature linked to CH_4_intensity, comparable to microbial traits previously associated with residual feed intake (RFI) or milk-fat phenotypes [45, 46]. Our results therefore reveal a microbial signature that extends beyond taxonomy, integrating digestive and metabolic drivers that differentiate CH_4_ intensity phenotypes.

**Figure 6.**
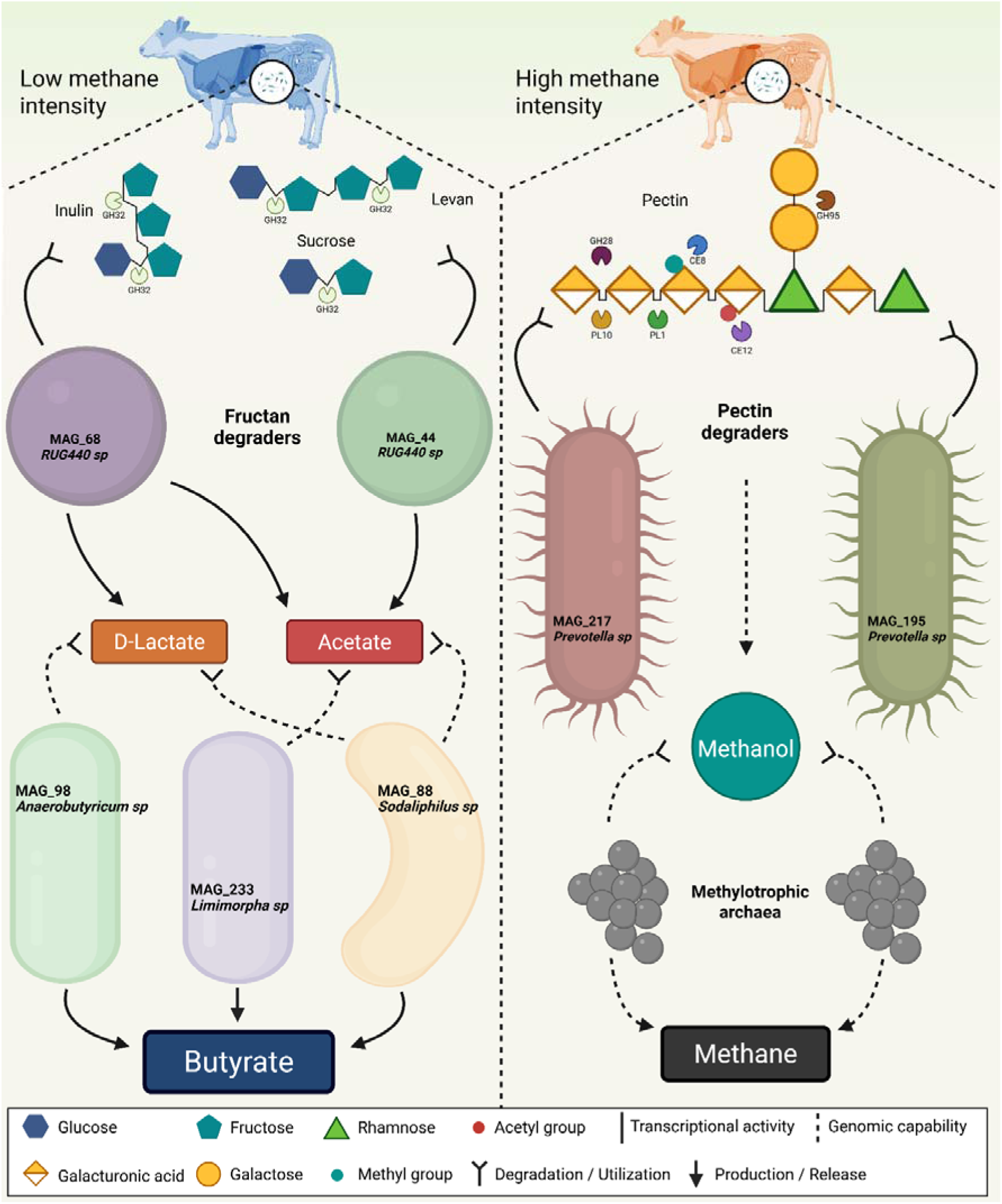
Reconstruction of carbohydrate degradation in low and high CH_4_ intensity cows. MAGs present showed a tendency between groups. Solid arrows are processes based on transcriptional activity, whereas dashed arrows are predictions based on genomic capability of MAGs. Created in BioRender. Maynez Perez, A. (https://BioRender.com/5mqp8w1).

The enriched expression of fructan-degrading genes in the LMI rumen indicates a microbial capacity to exploit this carbohydrate polymer, abundant in winter forages such as ryegrass (*Lolium perenne*) [47] and timothy (*Phleum pratense*) [48], which together contributed over 80% of dietary dry matter. Three RUG440 (*Atopobiaceae*) MAGs were identified as major fructan degraders, suggesting their ecological role as primary degraders within the rumen. Members of this genus have been reported in rumen environments dominated by *Epidinium spp.* [49], although their metabolic function remained unresolved. Additional MAGs encoding fructan-degrading genes belonged to *Sodaliphilus* and *Anaerobutyricum*, though with less transcript abundance than the RUG440 representatives. Their relevance lies in the genetic potential to convert metabolites produced by fructan degraders, such as lactate and acetate, into butyrate, together with *Limimorpha* (Fig. 6). These results suggest a possible cross-feeding relationship between fructan degraders and butyrate producers in the LMI rumen. Similar syntrophic interactions between lactate/acetate producers and butyrate formers have been documented for *Bifidobacterium*, *Lactobacillus*, and *Megasphaera* species [50–52], while the absence of lactate-to-butyrate converters has been reported in high-emitting cattle [53]. Consistent with this mechanism, rumen simulation studies supplementing inulin as a fructan source also observed increased butyrate production [50]. Although the influence of butyrate on milk yield remains uncertain, probiotic supplementation with native ruminal butyrate-producing strains has been shown to enhance milk yield in dairy cows [54].

Conversely, the HMI rumen showed enriched transcription of genes for degrading pectin, a polysaccharide ubiquitous in the primary cell wall of forage plants [55], including those in the experimental diet. Pectin structure involves the methyl-esterification of galacturonic acid residues [56]; hydrolysis of these methyl esters releases methanol, which can be stoichiometrically converted to CH_4_ (4 CH_3_OH → 3 CH_4_ + CO_2_ + 2 H_2_O) [57]. Although total CH_4_ production did not differ, all methanogenesis pathways exhibited higher transcript levels in the HMI phenotype. Mapping of pectin-degrading CAZymes to *Prevotella* MAGs aligns with the established saccharolytic capacity of this genus to utilize xylans and pectins for growth [58]. This aligns with evidence that *Prevotella* is the dominant rumen genus capable of converting methoxylated pectin into methanol, highlighting its potential role in supplying substrates for methylotrophic methanogenesis [59]. *Prevotella* is part of the ruminal core microbiome and the genera with the highest number of strains across it [60, 61]. This genus has been reported in animals classified as low CH_4_ emitters [62], though in those studies categorization was driven by differences in dry-matter intake rather than CH_4_ production. Other multi-omic analyses have linked *Prevotella*-enriched protein modules negatively with feed efficiency in beef cattle [63] and associated this genus with lipopolysaccharide biosynthesis and inflammation risk [64]. Given its dominance in the rumen ecosystem, a more detailed understanding of *Prevotella* function at a higher species/strain-level resolution across wider ecological niches is essential.

Collectively, the LMI phenotype was supported by a microbial ecosystem engaged in fructan degradation and butyrate formation, potentially via cross-feeding interactions. In contrast the HMI phenotype was characterized by an enhanced capacity for methyl-esterified pectin degradation, suspected of supplying methylotrophic archaea with methanol and supporting methanogenesis (Fig. 6). These findings provide insights into ruminal microbial fermentation and ecosystem function in relation to a less-studied CH_4_ metric (CH_4_ per ECM). Future work should involve larger cohorts and direct metabolite measurements (SCFA profiles, metaproteomics, and metabolomics) to validate these metagenomic and metatranscriptomic observations. Further, isolation of relevant microorganisms—particularly those capable of fructan degradation and butyrate production—combined with *in vitro* assays could clarify potential microbial cross-feeding relationships. Understanding the ruminal microbiome in efficient animals which produce less CH_4_ per unit milk could improve breeding programs and the development of microbial-based CH_4_-mitigation strategies.

## Acknowledgments

The authors gratefully acknowledge the staff at the Livestock Production Research Center (SHF) and the Metabolism Department (SSA) of the Norwegian University of Life Sciences for their expert animal care and invaluable assistance during sample collection. Special thanks are extended to TINE for formulating the experimental diet, and to DNASense for DNA and RNA sequencing. Computational resources were provided by the Orion High Performance Computing Center at the Norwegian University of Life Sciences and by Sigma2, the National Infrastructure for High Performance Computing and Data Storage in Norway. The authors also acknowledge ELIXIR Norway, supported by the Research Council of Norway (NFR grant 322392), for bioinformatics and data management support.

## Author contributions

AOMP: data curation, formal analysis of rumen multi-omics, investigation, software, visualization, writing – original draft; HNC: data curation, formal analysis of animal performance data, investigation, visualization, writing – review and editing; PN: data curation, writing – review and editing; VTEA: supervision, writing – review and editing, PBP: supervision, writing – review and editing; AS: conceptualization, supervision, project administration, funding acquisition, writing – review and editing.

## Competing interests

Author PBP has stock and/or equity interests in Bovotica Pty Ltd. The remaining authors declare no competing interests.

## Funding

This work was funded by the Research Council of Norway (ViableCow, project No 316157). PBP also acknowledges support from the Australian Research Council (Future Fellowship: FT230100560).

## Data availability

Metagenomic and metatranscriptomic sequences can be accessed at the European Nuleotide Archive (ENA, project number PRJEB105185).

