## Supplement for "Intricate microbiome differences observed in lactating cows across methane intensity phenotypes"

**Short title:** Fructan–butyrate in low-CH₄ intensity

Adrian Omar Maynez-Perez^1,2†^, Hendra Nur Cahyo^1,3†^, Puchun Niu^1,4^, Velma T. E. Aho^1^, Phillip B. Pope^1,5,6*^, Angela Schwarm^1*^

^†^These authors contributed equally to this work

^1^Department of Animal and Aquacultural Sciences, Norwegian University of Life Sciences, PO Box 5003, 1432 Ås, Norway

^2^Facultad de Zootecnia y Ecologia, Universidad Autonoma de Chihuahua, Chihuahua 31453, Chihuahua, Mexico

^3^Department of Animal Nutrition and Feed Science, Faculty of Animal Science, Universitas Gadjah Mada, 55281 Yogyakarta, Indonesia

^4^Department of Animal Science, College of Agriculture and Life Sciences, Cornell University, 215 Garden Ave, Ithaca, NY 14850, United States of America

^5^The Centre for Microbiome Research, Queensland University of Technology, 4102 Woolloongabba, Australia

^6^Faculty of Chemistry, Biotechnology and Food Science, Norwegian University of Life Sciences (NMBU), 1432 Ås, Norway

*Senior authors: Phillip B. Pope, Angela Schwarm

**Content**

Nonstandard abbreviations

Supplementary Materials and Methods

Supplementary Figures

Supplementary Tables

**Nonstandard abbreviations**

DM = dry matter; OM = organic matter; NDFom = ash-free neutral detergent fibre; ADFom = ash-free acid detergent fibre; CP = crude protein; NFC = non-fibre carbohydrates; ECM = energy corrected milk; BW = body weight.

**Supplementary Materials and Methods**

***Experimental setup and diet composition***

The trial was conducted at the Livestock Production Research Center (Senter for Husdyrforsøk, SHF) and the Metabolism Department (Stoffskifteavdeling, SSA), Norwegian University of Life Sciences. The experiment included two blocks of eight cows due to barn capacity. During adaptation, cows were housed for seven days at SHF and four days at SSA; all data collection occurred at SSA. Nine cows were primiparous (LMI = 5; HMI = 4) whereas seven where multiparous (LMI = 3; HMI = 4).

The diet consisted of grass silage and concentrate at an average silage:concentrate ratio of 83:17 (DM basis). The silage was prepared from second-cut ryegrass (*Lolium perenne*) and first-cut timothy mixture, the latter containing ~50% timothy (*Phleum pratense*), 20% meadow fescue (*Festuca pratensis*), 20% meadow grass (*Poa pratensis*), and 10% white clover (*Trifolium repens*). The concentrate (Drøve Fase 1; Norgesfôr, Norway) was formulated using TINE OptiFôr software based on average milk yield. Mineral–vitamin mix (Normin Inc., Norway) was provided at 100 g d⁻¹. Silage was offered three times daily (07:15, 12:30, 18:15 h) with concentrate fed 30 min after each silage meal and concentrate allowance through GreenFeed system (1.5 kg per day). Cows had free access to water throughout.

***Feed, faeces, and urine collection and analyses***

For chemical analyses, representative samples were collected and stored at –20 °C. Feed samples of ryegrass, timothy mix and concentrate were pooled each and freeze-dried for 4 days to constant weight, ground (1 mm sieve; Retsch ZM 100 or SM 200), and stored at room temperature until analysis.

During the 72-h total collection phase, faeces and urine were collected into separate polypropylene containers positioned behind each cow. Urine was stabilized with 10% sulfuric acid (final pH < 4) to prevent nitrogen loss. Total daily output was weighed, and 10% representative samples were pooled per cow and stored at –20 °C. Faecal pools were freeze-dried and ground for later nutrient analysis.

Dry matter (DM) was measured by oven-drying subsamples for 24 h at 103°C. Crude protein was calculated as total nitrogen × 6.25, with nitrogen determined using a Kjeltec 8400 analyser (Foss, Hillerød, Denmark) following AOAC method 2001.11 (AOAC International, 2002). Neutral detergent fibre (NDFom) and acid detergent fibre (ADFom) concentrations were quantified using an Ankom200 Fiber Analyzer (Ankom Technology, Macedon, NY). Chemical composition of the diet consumed by LMI and HMI cows was calculated (Table S1).

***Measurement of enteric gas***

Methane (CH₄) and carbon dioxide (CO₂) were measured using a portable open-circuit head chamber (GreenFeed, C-Lock Inc.). Each cow was sampled manually at staggered 5–6 h intervals for 6 days to cover a full diurnal cycle. Airflow was maintained above 26 L s⁻¹ (mean ± SD = 35.8 ± 4.6 L s⁻¹), and filters were cleaned routinely. Instrument calibrations (20% O₂ in N₂) were performed daily, and CO₂ recovery tests before, between, and after blocks. Each cow remained in the hood for ~5 min while receiving a 45 g bait every 30 s (≈ 1.5 kg DM d⁻¹). Background air was measured between sessions for baseline correction. For ensuring a coverage of methane emission across the day, 22 measurements per cow were performed spanning nearly the full 24-hour cycle.

***Milk yield and composition***

Milk yield was recorded twice daily (08:00–09:00 h; 19:00–20:00 h) using a Tru-Test Milk Meter. Forty-mL samples were preserved with bronopol tablets and stored at 4 °C until analysis (TINE Laboratory, Heimdal, Norway). Milk composition (fat, protein, lactose) was determined by infrared spectroscopy, and ECM was computed from yields weighted by milking time.

***Unbinned-read taxonomic profiling***

To assess consistency in diversity patterns between the assembled and unassembled fractions, taxonomic profiling of raw metagenomic reads was performed using Kraken2 (v2.1.2) [1] with the PlusPF-8 database. Species-level abundance estimates were refined using Bracken (v2.9) [2] with parameters adjusted for long-read sequencing (read length = 1000 bp) (Figure S2).

**Supplementary Figures**

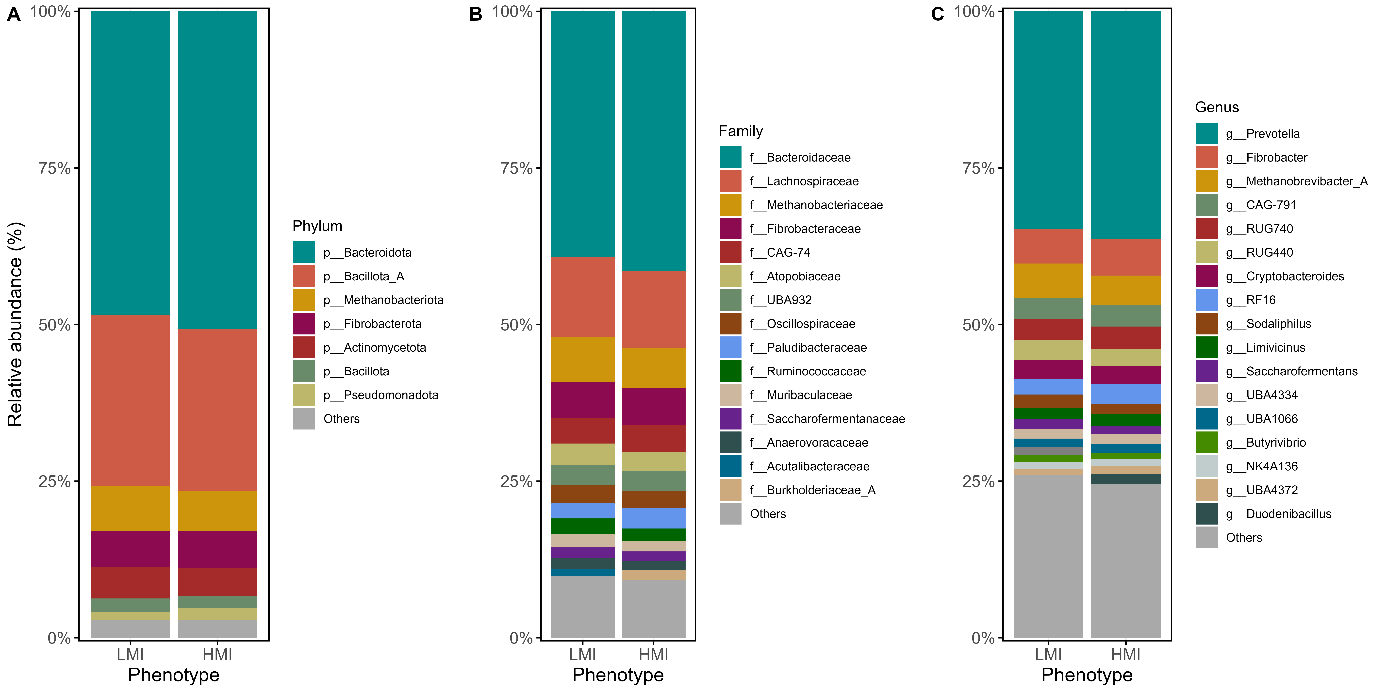

***Figure S1.*** *Microbial relative abundance in low (LMI) and high (HMI) CH_4_ intensity cows at the level of phylum (A), family (B), and genus (C); microbial groups with a relative abundance <1% are labelled as “Others”.*

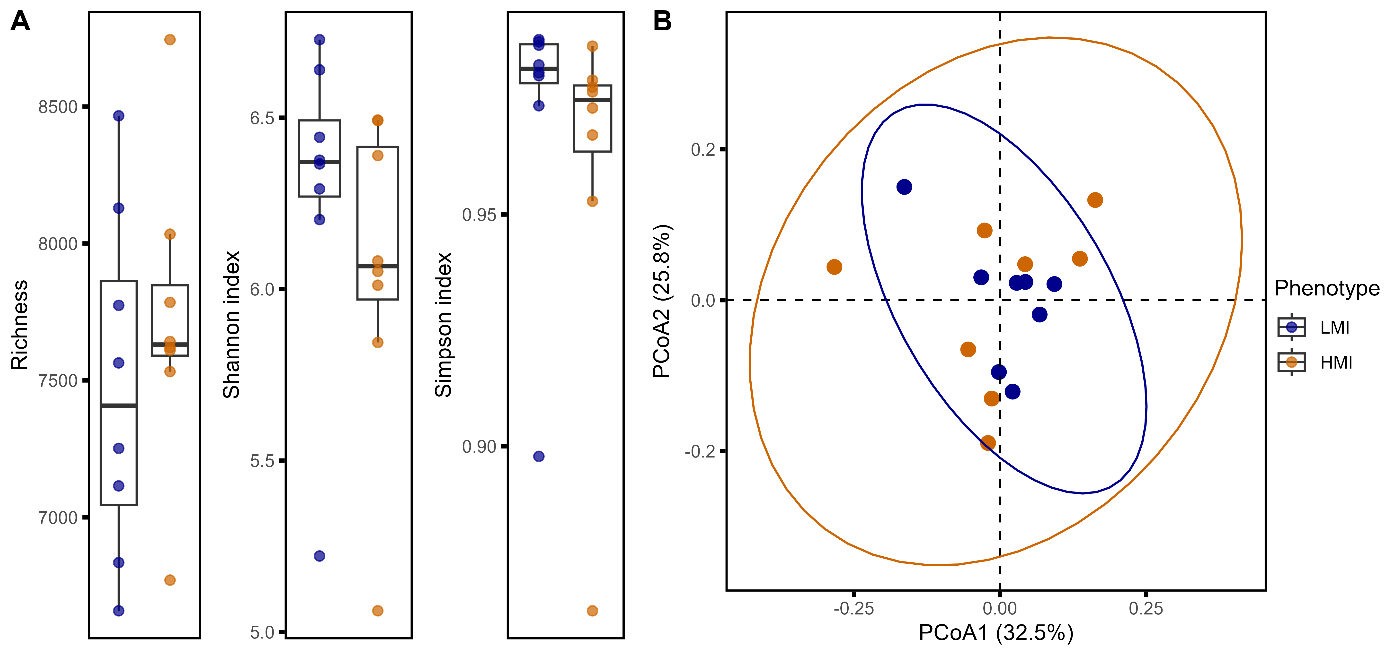

***Figure S2.*** **Alpha and beta diversity of the unbinned metagenomic fraction in rumen samples from low (LMI) and high (HMI) CH_4_ intensity cows**. ***A.*** *Boxplots displaying alpha diversity including richness, Shannon and Simpson index. No difference was found in any of the metrics (FDR = 0.50).* ***B.*** *Principal coordinates analysis (PCoA) plot illustrating beta diversity based on Bray-Curtis method. No difference was found between phenotypes (FDR = 0.81).*

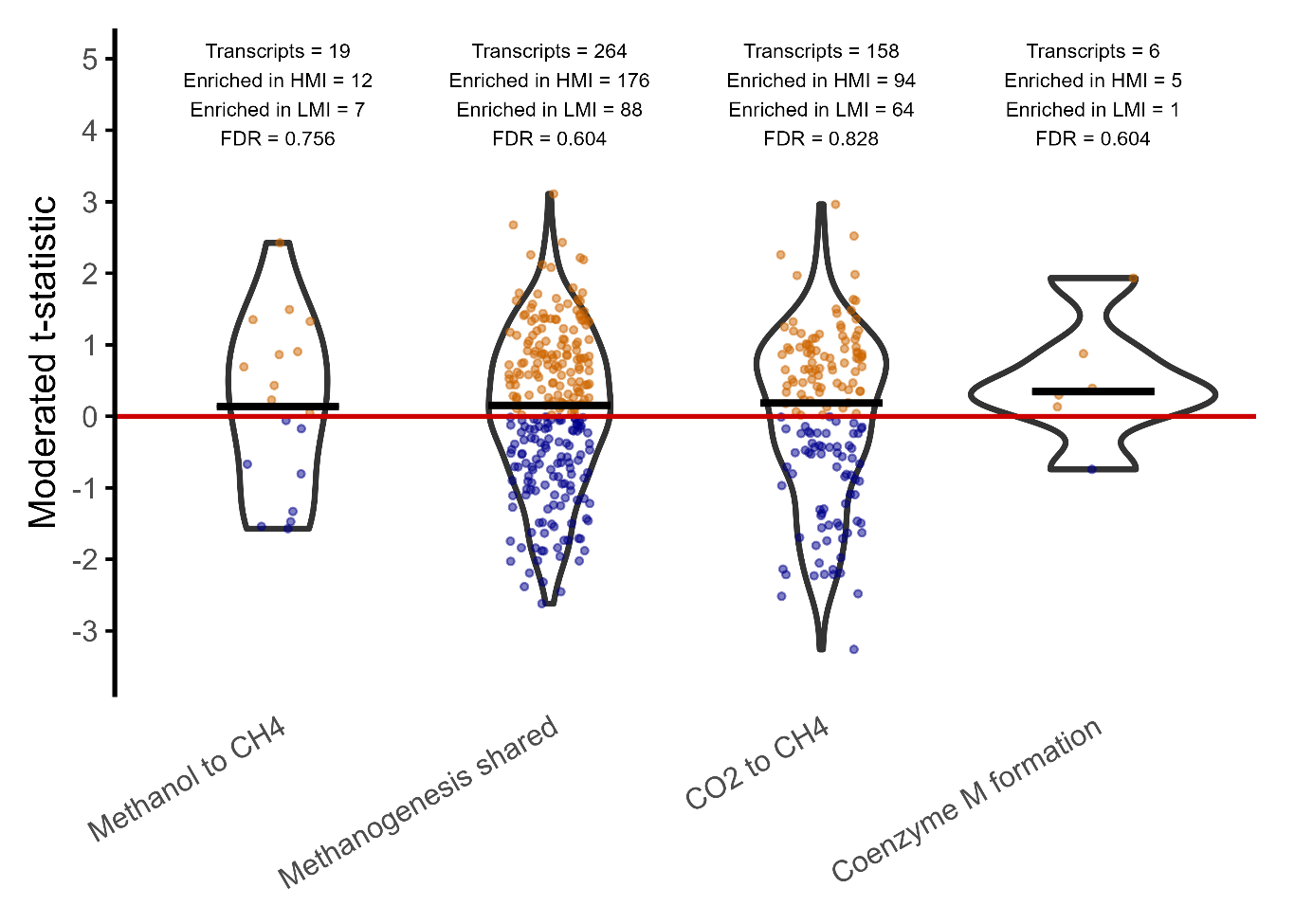

***Figure S3.*** *Comparative* *metatranscriptomic analysis of rumen methane pathways* (KEGG)*. Each dot (gene) represents the moderated t-statistic from a limma–voom model contrasting phenotypes (HMI = high CH_4_ intensity vs LMI = low CH_4_ intensity). Outlines depict the kernel density of gene-level t-statistics per set, whereas the black crossbar marks the median. The colour of each dot encodes direction where positive values (orange) indicate higher expression in the HMI group, whereas negative values (blue) indicate higher expression in the LMI group. Statistical significance was determined by CAMERA gene-set testing applied to the voom (log-CPM) matrix, adjusted by FDR (Benjamini–Hochberg).*

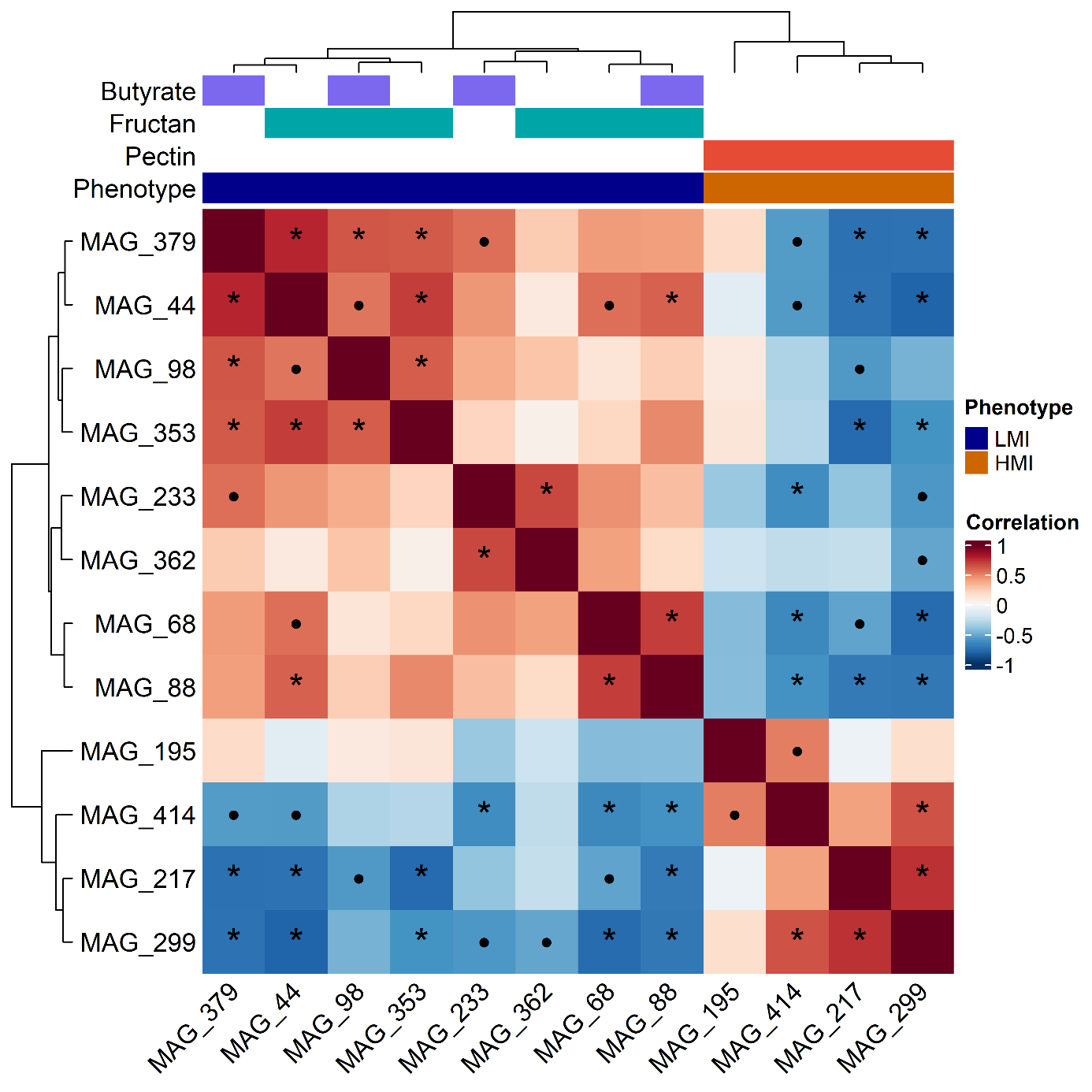

***Figure S4.*** *MAG–MAG association structure after compositional adjustment. Heatmap shows pairwise Spearman correlation (ρ) between abundances of MAGs encoding pathways of interest. Data was centred log-ratio (CLR) transform. Rows and columns are hierarchically clustered. Significance marks inside cells denote multiple-testing–adjusted support (Benjamini–Hochberg across all pairs): FDR< 0.05 (*), FDR < 0.10 (•). Top annotation tracks indicate guild membership (butyrate metabolism, fructan degradation, and pectin degradation; a MAG can show multiple tracks) and CH_4_ intensity phenotype (LMI or HMI).*

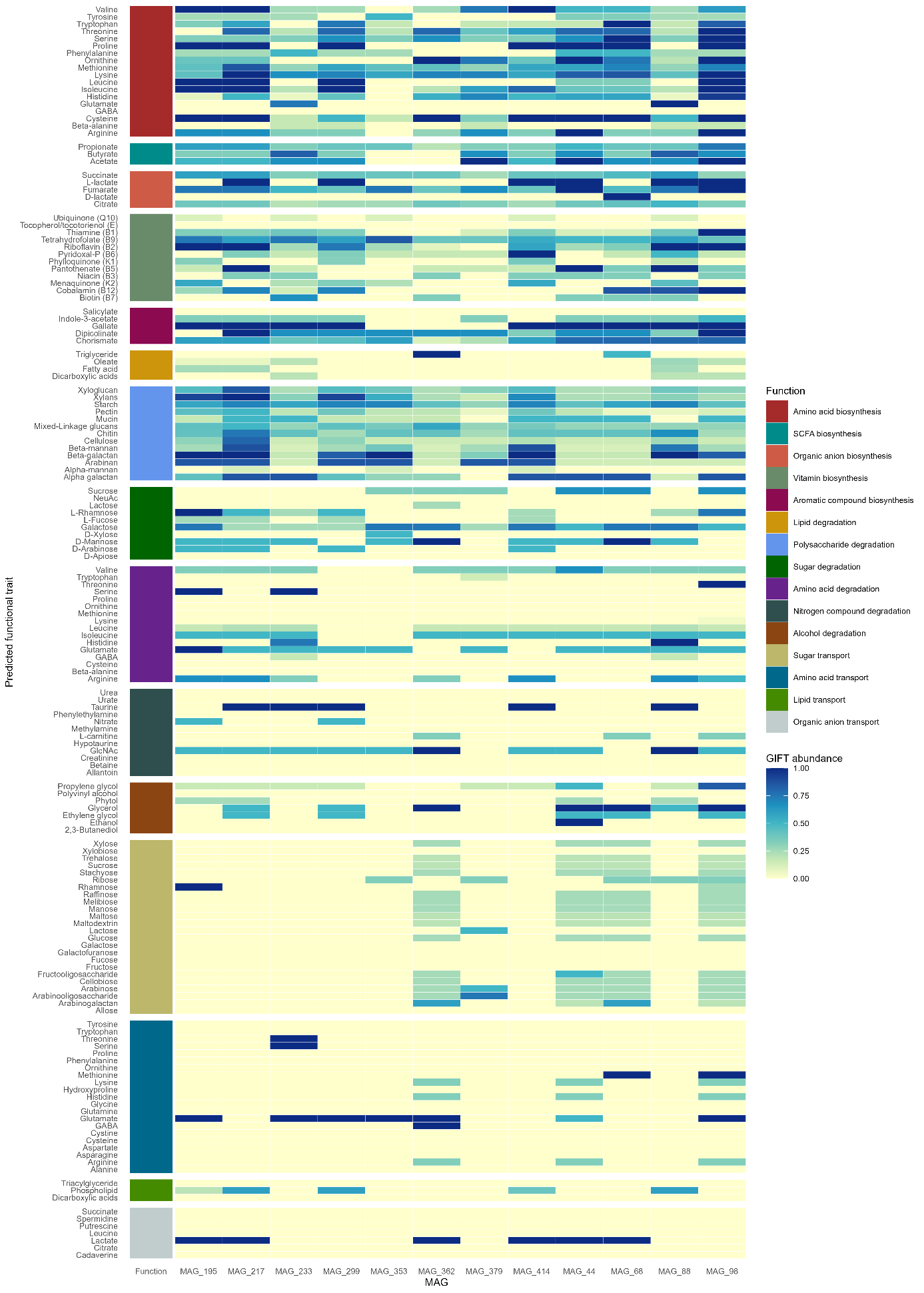

***Figure S5.*** *Genome-inferred functional traits (GIFTs) across tendency MAGs. Heatmap shows the abundance of genome-inferred functional traits linked to targeted metabolic functions for MAGs that exhibited a directional tendency (non-adjusted P < 0.05 from a two-sided Wilcoxon rank-sum test) with CH_4_ intensity (LMI or HMI).*

**Supplementary Tables**

***Table S1****. Chemical composition (% of DM, unless stated otherwise) of the diet consumed by low (LMI) and high methane intensity cows (HMI)*

| Item | Diet^1^ consumed | |
| --- | --- | --- |
|  | LMI | HMI |
| DM, % of fresh matter | 34.9 | 35.0 |
| OM | 91.6 | 91.6 |
| Ether extract | 3.18 | 3.18 |
| NDFom | 46.1 | 46.0 |
| ADFom | 28.3 | 28.2 |
| CP | 16.7 | 16.8 |
| NFC^2^ | 24.6 | 24.7 |
| Gross energy, MJ/kg of DM | 19.1 | 19.1 |

^1^The diet consisted of grass silage and concentrate with an average ratio of 83:17 on a DM basis. Grass silage comprised of a mixture of two bales of ryegrass (*Lolium perenne*) and one bale of a mixture of timothy grass (50%, *Phleum pratense*), meadow fescue (20%, *Festuca pratensis*), meadow grass (20%, *Poa pratensis*), and white clover (10%, *Trifolium repens*) in the seed mixture (fresh matter basis). The energy concentrate (Drøv Fase 1, Norgesfôr, Norway) comprised pelleted barley, 47.6%; SoyPass (soy-based bypass protein), 12.2%; unmolassed beet pulp, 7.5%; white field beans (not heat treated), 5.56%; rapeseed expeller, 5%; maize, 5%; maize gluten, 3%; pelleted oats 2.75%, Lipitec Bovi LM, 2.73%; salt, 0.87%; monocalcium phosphate, 0.66%; fine calcite, 0.33%;  Trouw levucell premix, 0.25%; Vilomix drøv, 0.25%; calseapowder advance, 0.70%; magnesium oxide, 0.6%. Vitamin and mineral content per kg concentrate product: vitamin A, 4000 IU; vitamin D3, 2000 IU; vitamin E, 60 IU; Mn, 30 mg; Cu, 15 mg; Zn, 70 mg; I, 5 mg; Co, 0.4 mg; Se (sodium selenite), 0.3 mg; Se (selenium-enriched inactivate yeast) 0.1 mg, and live yeast (*Saccharomyces cerevisiae*) at level 1 billion CFU per kg of product.

*^2^Non-fiber carbohydrates (NFC): 100-(NDF+CP+EE+Ash); Ash: 100-OM.*

***Table S2****. Nutrient intake, total tract digestibility, milk yield and composition, methane and carbon dioxide production, energy intake and loss in cows with low (LMI) and high CH_4_ intensity (HMI).*

| Item | Phenotype | | SEM | FDR |
| --- | --- | --- | --- | --- |
|  | LMI | HMI |  |  |
| Nutrient intake, kg/d |  |  |  |  |
| DM | 20.0 | 18.7 | 0.963 | 0.38 |
| OM | 17.5 | 16.4 | 0.790 | 0.35 |
| Apparent digestibility, % | |  |  |  |
| DM | 76.6 | 75.4 | 1.13 | 0.48 |
| OM | 77.5 | 76.4 | 1.09 | 0.50 |
| Milk yield, kg/d | 20.1 | 14.4 | 0.938 | <0.01 |
| ECM, kg/d | 20.9 | 15.5 | 0.995 | <0.01 |
| ECM/DMI, kg/kg | 1.10 | 0.86 | 0.031 | <0.01 |
| Milk composition | |  |  |  |
| Fat, % | 4.22 | 4.44 | 0.168 | 0.38 |
| Protein, % | 3.65 | 3.84 | 0.108 | 0.24 |
| Lactose, % | 4.48 | 4.55 | 0.064 | 0.46 |
| CH_4_ |  |  |  |  |
| g/d | 431 | 419 | 15.1 | 0.60 |
| g/kg of DMI | 22.6 | 23.6 | 0.672 | 0.33 |
| g/kg of BW^0.75^ | 3.43 | 3.30 | 0.121 | 0.45 |
| g/kg of ECM | 20.7 | 27.6 | 1.12 | <0.01 |
| CO_2_ |  |  |  |  |
| g/d | 12227 | 12076 | 443 | 0.81 |
| g/kg of DMI | 641 | 678 | 13.9 | 0.09 |
| g/kg of BW^0.75^ | 97.3 | 94.7 | 2.20 | 0.43 |
| g/kg of ECM | 587 | 794 | 28.4 | <0.01 |
| Energy intake, MJ/d |  |  |  |  |
| Gross energy (GE) | 369 | 346 | 16.7 | 0.35 |
| Digestible energy | 274 | 254 | 14.6 | 0.34 |
| Metabolizable energy | 232 | 214 | 14.2 | 0.39 |
| Energy loss, MJ/d |  |  |  |  |
| Methane^1^ | 23.7 | 23.1 | 0.818 | 0.63 |
| Milk energy | 65.7 | 48.5 | 3.13 | <0.01 |

^1^CH_4_ energy (MJ/d) = CH_4_ (L/d) × 0.03957. The conversion factor of 0.7168 was used to assume the density of CH_4_ gas in gram per Liter at standard temperature and pressure.
